# The lonely fish is not a loner fish: whole-brain mapping reveals abnormal activity in socially isolated zebrafish

**DOI:** 10.1101/2020.01.22.915520

**Authors:** Hande Tunbak, Mireya Vazquez-Prada, Thomas Ryan, Adam R. Kampff, Elena Dreosti

## Abstract

The zebrafish is used to assess the impact of social isolation on behaviour and brain function. As in humans and other social species, early social deprivation reduces social preference in juvenile zebrafish. Whole-brain functional maps of anti-social isolated fish were distinct from anti-social fish found in the normal population. These isolation-induced activity changes revealed profound disruption of neural activity in brain areas linked to social behaviour, such as the preoptic area and hypothalamus. Several of these affected regions are modulated by serotonin, and we found that social preference in isolated fish could be rescued by acutely reducing serotonin levels.

## Short Report

Social preference behaviour (Rogers-Carter et al., 2018; Winslow, 2003) the drive for individuals to identify and approach members of their own species, is a fundamental component of all social behaviour. We previously found that most zebrafish develop a strong social preference by 2-3 weeks of age (Dreosti, Lopes, Kampff, & Wilson, 2015), yet a small portion of the population (∼10%) was averse to social cues. Similar diversity in individual preferences have been found in populations of many social species, including humans (Sloan Wilson, Clark, Coleman, & Dearstyne, 1994). Isolation from social interaction has been linked to a reduction in social preference (Engeszer, Ryan, & Parichy, 2004; S Shams, Amlani, Buske, Chatterjee, & Gerlai, 2018).Therefore, we first asked whether isolation would impact the behaviour of zebrafish in our visual social preference assay.

Fish were placed in full isolation from fertilization to 3 weeks, as described in the Methods section, and then tested in the behavioural assay. Each experiment comprised two sessions, 15 minutes of acclimation to the chamber followed by 15 minutes of exposure to two size matched sibling fish, which were not isolated. To quantify social preference, we calculated a Visual Preference Index (VPI) that compares the amount of time fish spend in the chamber nearest the conspecifics versus the opposite chamber where they are visually isolated from social cues (see Methods). Full social isolation (Fi) caused a significant decrease in social preference relative to normally raised sibling controls (C) (Figure 1A, left and middle panel: C vs Fi, p= 8.3 e^−8^, Mann-Whitney). Specifically, there was an increase in the number of individuals that had a large negative VPI. We, therefore, decided to divide the fish into three groups: a) anti-social (-S) fish with VPIs below -0.5; b) pro-social (+S) fish with VPIs above +0.5; c) non-social (ns) fish with -0.5 < VPI < +0.5. Fish that underwent Partial isolation (Pi), i.e. isolated from social cues for 48 hrs immediately prior to the behavioural experiment, exhibited an intermediate, yet highly significant, change in social preference (Fig.1A, right panel, C vs Pi, p=2.5e^−8^, Mann-Whitney).

**Figure 1.**
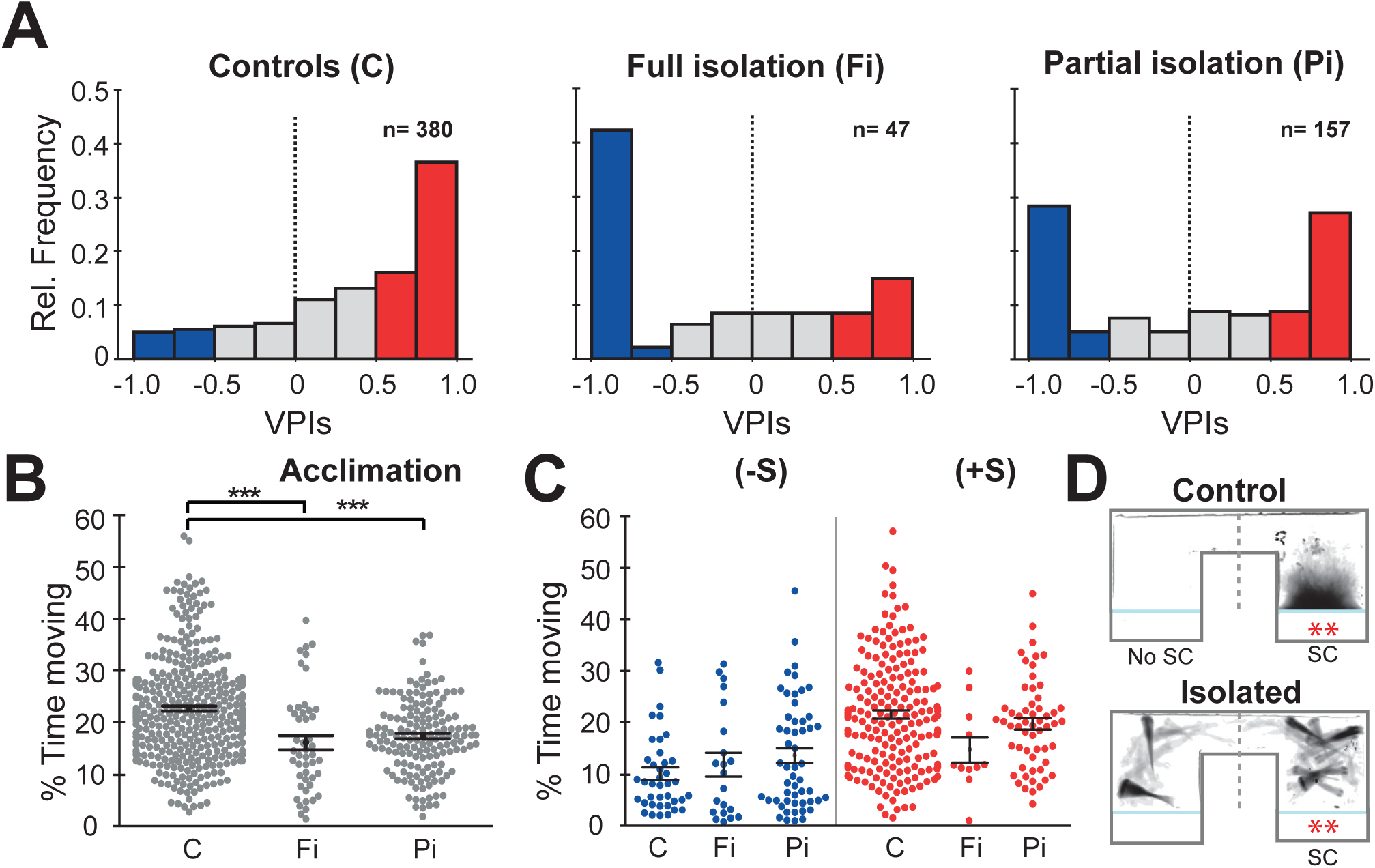
Isolation alters social preference behavior and swimming activity. **A**. Histograms of all the VPIs during the social cue period across different conditions: controls (C, left), full isolation (Fi, middle), and partial isolation (Pi, right). For visual clarity, red bars highlight strong pro-social fish (+S, VPIs > 0.5), blue bars anti-social fish(-S, VPIs < -0.5), and gray non-social fish (ns, -0.5 < VPI < +0.5). **B**. Swarm plots comparing the activity levels of fish during the acclimation period expressed as percent time moving (C, n=380; Fi, n=47; Pi, n=157). Mean and standard errors are shown. **C**. Swarm plots comparing the activity levels of anti-social (left) and social (fish) fish during visual social cue exposure for each rearing condition (C (-S), n=39; Fi (-S), n=21; Pi (-S), n=53) or (C (+S), n=193; Fi, n=11; Pi (+S), n=57). **D**. Time projection through the video of a pro-social control, C(+S), and a fully isolated, Fi(+S), fish during social cue exposure. The dashed marks the division between the social cue side (SC) and the side without social cues (No SC) that was used to calculate VPI.

As previously reported (Zellner, Padnos, Hunter, MacPhail, & Padilla, 2011), we found that fish raised in full isolation were significantly less active than their normally raised siblings during the acclimation session (Figure. 1B, C vs Fi, p=9.0e^−6^, C vs Pi, p=2.8e-^9^ Mann-Whitney). However, when we compared the swimming activity of anti-social isolated and anti-social control fish, there were no significant differences (Figure 1C; C (-S) vs Fi (-S), p=0.48, Mann-Whitney). We then compared the swimming behaviour of isolated and normally raised anti-social (-S) fish in more detail and noticed that fish from both groups exhibited prolonged periods of quiescence (freezing) (Figure 1C left, C (-S) vs Fi (-S), p=0.48, Mann-Whitney). This freezing behaviour is consistent with similar anxiety behaviours observed in many species, and previously reported in adult zebrafish exposed to stressors (Giacomini et al., 2015; S Shams, Seguin, Facciol, Chatterjee, & Gerlai, 2017). The pro-social fully isolated fish, Fi (+S), instead, showed a different behavioural phenotype compared to the pro-social normally raised fish, C (+S), exhibiting an increase in freezing behaviour especially when observing conspecifics (Figure 1C right, Figure 1D, and Supp. Movie 1).

Anxiety-like behavioural responses have been shown to increase following periods of social isolation (S Shams et al., 2017) and they are also known to vary across individuals in a population (Egan et al., 2009). The behavioural similarities between anti-social fish isolated and anti-social fish found in the normal population led us to hypothesize that maybe isolation predisposes fish to a high-anxiety state. If this is the case, neural activity of anti-social isolated and normally raised fish should be similar when presented with social cues. To test this hypothesis, we performed whole-brain two-photon imaging of *c-fos* expression, an immediate early gene whose expression is associated with increased neural activity(Herrera & Robertson, 1996), in juvenile brains following testing in the social preference assay. Dissected brains were imaged with the dorsal surface down (bottom-up) to achieve clear views of the ventral brain structures that have been previously implicated in the social brain network (Figure 2A, see Methods). Volumes of 1.5 mm x 1.5 mm x 700 µm, with a voxel size of 1×1×3 µm (2048×2048 pixels), were acquired from 135 zebrafish brains across all experimental groups and registered to a reference brain (Marquart et al., 2017).

**Figure 2.**
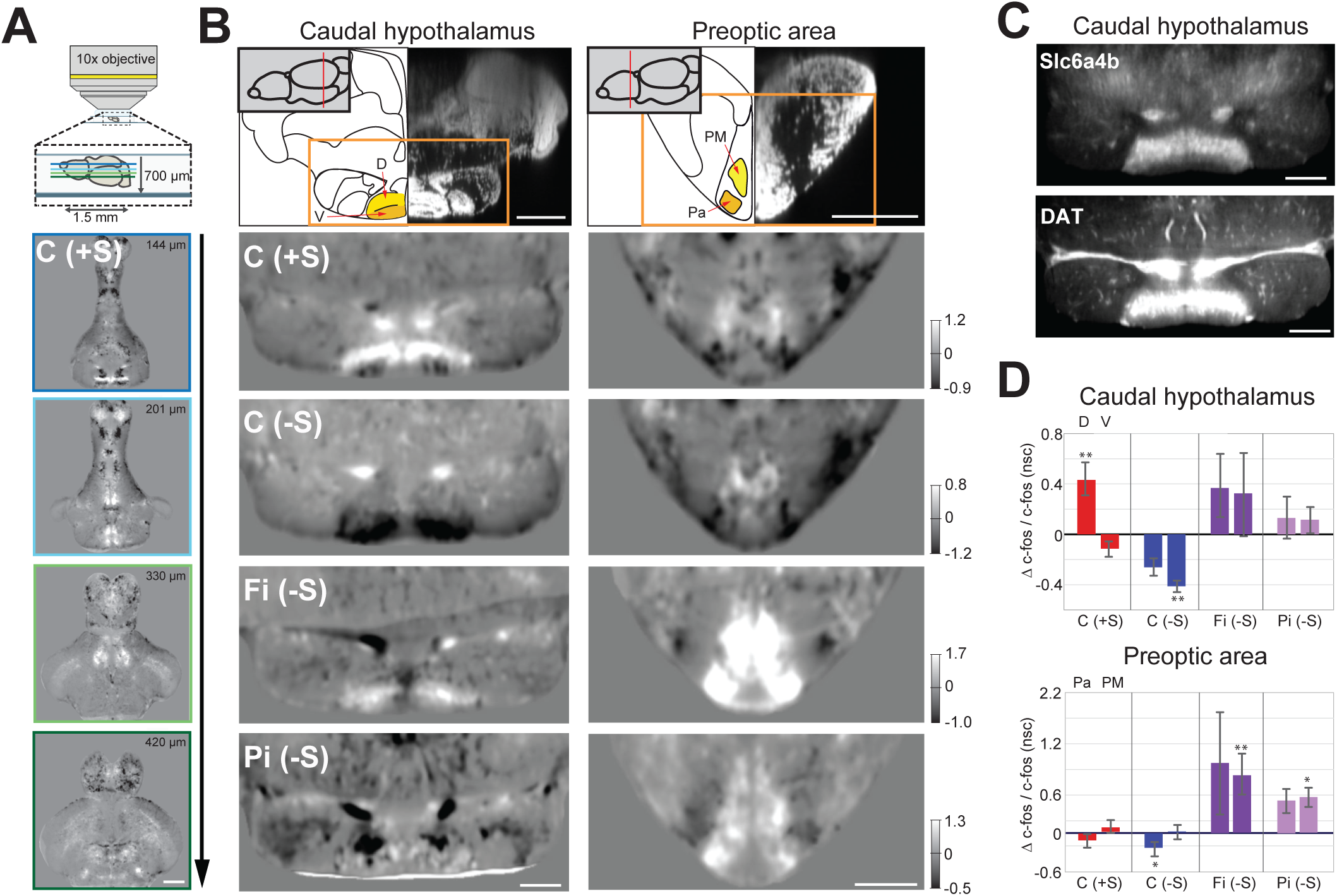
Functional maps of the social brain in normal and isolated fish. **A**. Schematic of the custom-built two-photon microscope used for acquiring whole-brain volumes of dorsal-down mounted fish brains (top panel). Horizontal sections, at increasing imaging depth, showing the average difference between C (+S) and siblings not presented with a social cue (lower panels). Positive values (white) indicate increased cFos expression in socially preferring fish, while negative values (black) indicate decreased expression. Scale bar is 200μm. The intensity scale bar is shown in Fig. 2B, C(+S) row. **B**. Region analysis of two different brain areas that have been implicated in social behavior: caudal hypothalamus and preoptic area. A schematic of the anatomical regions and corresponding DAPI staining is shown (top panel) with two sub-regions highlighted. Images showing changes in cFos activation in these areas for pro-social controls (C(+S)), anti-social controls (C(-S)), anti-social fully isolated (Fi(-S)), and anti-social partially isolated (Pi(-S)) are shown. Images are horizontal sections of the average difference between each test group and their corresponding sibling group not presented with a social cue. Scale bar is 100μm. Intensity scale bar is shown for each group. **C**. Average image of Slc6a4b and DAT expression in the same section of the caudal hypothalamus as 2B (n=3 each). Scale bar is 100μm. **D**. Summary graphs showing the change in cFos activation, relative to each group’s no social cue (nsc) siblings, for each sub-region of the social brain areas indicated in Fig. 2B (yellow/orange). Positive values indicate increases in cFos expression; asterisks mark significant changes relative to no social cue siblings. Pa=ventrolateral preoptic area, PM=dorsal preoptic area.

These *c-fos* whole-brain functional maps allowed us to compare the neural activity patterns of different test groups. For example, normally reared pro-social fish, C (+S) and a control group of sibling fish that were placed in the behavioural assay for 30 minutes without any social cue (no social-cue controls: (nsc)). The resulting average difference stacks (C (+S) vs. C (nsc)) were used to identify changes in neural activity associated with exposure to a visual social cue (Figure 2A).

We first investigated the regions activated in pro-social normally raised fish (Figure 2B C (+S)) that have been previously reported to be social brain areas (O’Connell & Hofmann, 2011). Coronal sections are shown for two different areas: caudal hypothalamus and the preoptic area. Significant activation in a region of the dorsal caudal hypothalamus was a prominent signature of C (+S) fish (Figure 2B and 2D, C(+S) vs C (nsc) p=0.007, Mann-Whitney). In contrast, anti-social, C (-S), fish showed a large decrease in activation in an adjacent, non-overlapping area of the ventral caudal hypothalamus (Figure 2B and 2D, C (-S) vs C (nsc) p=0.003, Mann-Whitney test). This functional division in the hypothalamus seems to mirror the distribution of dopaminergic and serotoninergic markers (Filippi, Mahler, Schweitzer, & Driever, 2010; Lillesaar, 2011) (Figure 2C), suggesting that this area could be responsible for the changes of serotonin and dopamine levels widely reported in fish viewing social cues (Huang et al., 2015; Soaleha Shams, Amlani, Buske, Chatterjee, & Gerlai, 2018). In the preoptic area, we observed only a small dorsal (PM) increase and ventrolateral (Pa) decrease in activity, which was significant only in the Pa of anti-social control fish (C(-S) vs C (nsc) p=0.003 Mann-Whitney, Figure 2B and 2D). These results are consistent with brain areas that have been previously identified as involved in social behaviour in a number of species (O’Connell & Hofmann, 2011).

We were now able to ask whether neural activity was similar in anti-social isolated and anti-social normally raised fish following the presentation of a social cue. Contrary to our hypothesis, *c-fos* functional maps of anti-social fully isolated fish (Fi (-S), Figure 2B) revealed a completely different activity profile than their anti-social normally raised siblings (C (-S), Figure 2B). The ventral caudal hypothalamus of Fi (-S) fish was not inactivated, and the preoptic was strongly activated in both dorsal (PM) and ventral (Pa) regions, but significantly only in the PM (Figure 2B and 2D, Fi (-S) vs Fi (nsc) p=0.006 PM; p=0.07 Pa, Mann-Whitney). Similar functional activity changes were observed in fish exposed to a brief isolation of only 48 hours prior to testing. (Pi (-S), Figure 2B), albeit less strong (Pi (-S) vs Pi (nsc) p=0.04 Pa; p=0.04 PM, Mann-Whitney, Figure 2D). The altered functional brain maps of isolated fish provide strong evidence that social isolation, while reducing social preference behaviour, does not do so by the same mechanism that produces social aversion in normally reared fish.

We were also interested in understanding why social isolation promotes social aversion instead of increasing the drive towards social cues. An important clue was found in the pattern of brain changes that were unique to isolated fish. When we compared activity levels in the functional brain maps of fully isolated fish (Fi (nsc); Figure 3A) which had never been exposed to social cues to normally raised controls, we found a significant increase in areas associated with visual processing (the optic tectum (McDowell, Dixon, Houchins, & Bilotta, 2004)) and stress responses (the posterior tuberal nucleus, PTN (Ziv et al., 2013).

**Figure 3.**
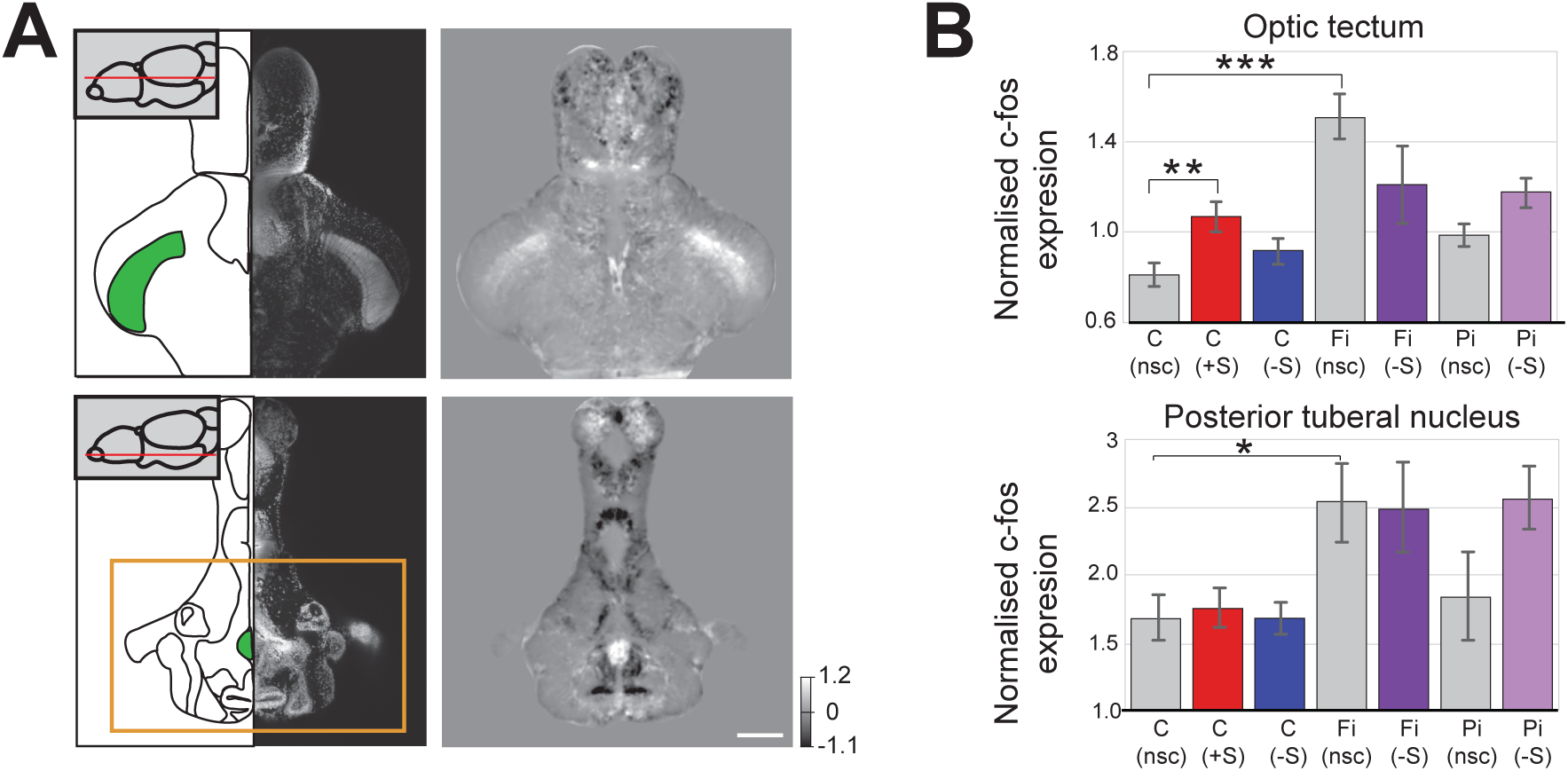
Changes in baseline brain activity following isolation. **A**. Images of two areas that show strong c-fos activation in fully isolated fish independent of social stimuli (optic tectum and posterior tuberal nucleus (PTN)). Schematics of the horizontal sections and corresponding DAPI image are shown in the left panels with the regions used for subsequent analysis indicated (green). Average images showing the difference in c-fos activation of fully isolated (Fi) and normally raised (C) fish without social cues (nsc) are shown in the right panels. Scale bar 200 μm. **B**. Summary graphs showing the normalised c-fos expression in the optic tectum and PTN regions indicated in Fig. 3A (green) for each experiment condition: normally raised controls without social cue C (nsc); pro-social controls C (+S); anti-social controls C (-S); anti-social fully isolated Fi (-S); fully isolated without social cue Fi (nsc); anti-social partially isolated Pi (-S); partially isolated without social cue Pi (nsc).

In pro-social control fish (C (+S), viewing a social cue causes a significant increase of neuronal activity in visual brain areas like the tectum (Figure 3B top; C (+S) vs C (nsc) p=0.004 Mann-Whitney). However, in fully isolated fish, there was already an increase in neuronal activity in the tectum in the absence of social cues (Fi (nsc)). This suggests that isolation increases visual sensitivity, as previously reported in humans (J T Cacioppo, Cacioppo, Capitanio, & Cole, 2015); (Figure 3B = Fi (nsc) vs C (nsc) p=0.0005 Mann-Whitney). This increased sensitivity of fully isolated fish (Fi (nsc)) was not found in partially isolated fish, Pi (nsc), suggesting that the visual sensitization effects of isolation are cumulative. In addition, increased tectal activity was also found in both full and partially isolated anti-social fish that were exposed to social cues (Fi(-S) and Pi(-S), even though these fish largely avoided the chamber with visual access to conspecifics (Fi (-S) vs C (nsc) p=0.016, and Pi (-S) vs C (nsc) p=0.0007, Mann-Whitney, Figure 3B).

A similar cumulative response was observed in the posterior tuberal nucleus of fully isolated fish. Full isolation caused a significant increase in PTN activity in absence of social cues (Fi (nsc) vs C (nsc) p=0.015, Mann-Whitney, Figure 3B bottom), and in anti-social fish (Fi(-S). During partial isolation, PTN activity was slightly increase and variable in absence of social cues (Pi(nsc), but significantly increased as well in anti-social fish (Pi(-S)). In humans, loneliness is associated with stronger activation of the visual cortex in response to negative social images (John T. Cacioppo, Norris, Decety, Monteleone, & Nusbaum, 2009) and it has been shown to increase attention to negative social stimuli (e.g. rejection, social threats, and exclusion). We therefore propose that social isolation in fish initially activates neuronal and neuroendocrine responses that promote anxiety-like behaviors, such as increased vigilance for social threats, hostility and social withdrawal, as observed in humans (John T. Cacioppo et al., 2009).

To test whether reducing anxiety could reverse the anti-social behaviour observed in isolated zebrafish, we acutely treated control and partially isolated fish with Buspirone, a drug that has been shown to reduce anxiety in humans, mice, and zebrafish (Bencan, Sledge, & Levin, 2009; Lalonde & Strazielle, 2009; Lau, Mathur, Gould, & Guo, 2011; Patel & Hillard, 2006) Buspirone, an agonist of the auto-inhibitory 5HT1A receptors, has been shown to enhance social interaction of rats (File & Seth, 2003; Gould et al., 2011) and sociability of zebrafish (Barba-Escobedo & Gould, 2012), and social phobia in humans (Schneier et al., 1993; van Vliet, den Boer, Westenberg, & Pian, 1997) Its ability to counter the effects of social isolation in zebrafish has not been investigated.

We first tested the effects of acute exposure to Buspirone in control fish, and, as expected, we observed a small significant increase in social preference relative to untreated controls (Supp. Figure 1A-B: C (no drug) vs C (30 µM) p=0.01 Mann-Whitney). We then treated partially isolated fish with 30 µM (Supp. Figure 1A-B, n=46 fish) and 50 µM (Supp Figure 1A-B, n=72 fish) Buspirone. Remarkably, the acute drug treatment was sufficient in both concentrations to reverse the anti-social phenotype caused by isolation (Figure 1A and 4A; Pi vs Pi (Buspirone 30 µM and 50 µM combined), p= 2.56 e-05 Mann-Whitney).

When we compared the time course of this phenotype reversal by computing the VPIs for each minute throughout the 15 minutes of the behavioural experiment (Figure 4 B), we found that the Buspirone treated isolated fish, while initially anti-social, would rapidly recover normal social preference behaviour within the first 5 minutes of exposure to social cues (VPI at 0 and 3 minutes between Pi and Pi(Buspirone) p= 0.01, and p=0.09 Mann-Whitney). In cotrast, the VPIs of untreated isolated fish remained significantly lower than controls throughout the entire session. Buspirone’s impact on the rate of recovery of social preference suggests it may be reducing anxiety by regulating circuit plasticity, perhaps by promoting down-regulation of the hypersensitivity acquired during the isolation period.

**Figure 4.**
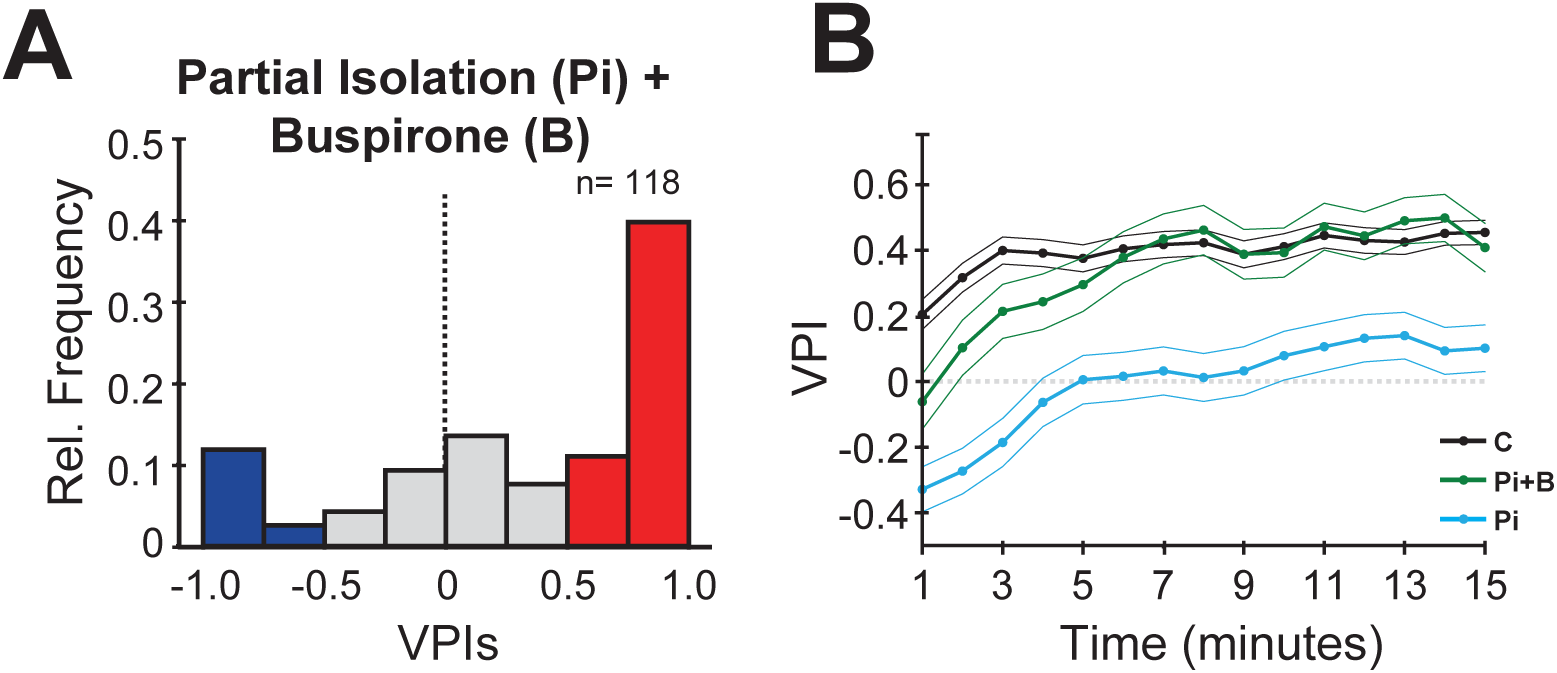
Buspirone rescues social preference in isolated fish. **A**. Histogram of VPIs during the social cue period in partially isolated (Pi) fish treated with 30 μM and 50 μM of Buspirone (combined). For visual clarity, the bars are colored as in Figure 1. **B**. VPI values calculated in one minute time bins for controls (C, black line, n=380), partial isolated (Pi, blue line, n=157), and Pi treated with Buspirone (Pi+B, green line, n=118). Note how Buspirone treated fish recover normal social preference within the first 5 minutes.

In summary, our results demonstrate that *lonely fish*, which have been isolated from social cues and show anti-social behaviour, have a completely different brain activity compared to *loner* anti-social fish found in the normal population. In addition, the functional changes caused by social deprivation are consistent with an increase in anxiety state found in humans, and could be reversed with an existing anxiolytic drug that acts on the monoaminergic system. Zebrafish, thus, will provide a powerful new tool for studying the impact of loneliness (isolation) on brain function and exploring different strategies for reducing, or even reversing, its effects.

## Methods

### Animals

AB strain zebrafish maintenance and breeding was performed at 28.5C with a 14h:10h light-dark cycle. Isolated fish were housed in custom chambers (length=15 cm, width=5 cm, height= 10 cm) made of opaque white acrylic with translucent lids, either from fertilization (full isolation) or for 48 hours prior to the behavioural experiment (partial isolation). All experiments were performed according to protocols approved by local ethical committee (AWERB Bloomsbury Campus UCL) and the UK Home Office.

### Behavioural assay and analysis

Experimental details and image acquisition were performed as described previously (Dreosti et al., 2015) and the source code can be found at http://www.dreo-sci.com/resources/. The visual preference index (VPI) was calculated by subtracting the number of frames in which the fish was located on the side of the arena nearest the social stimulus (Social cue (SC) side in Figure 1B) by the number of frames located on the opposite side of the arena (nsc (No SC) side). This difference was then divided by the total number of frames recorded [VPI = (SC – No SC)/Total frames].

### Whole mount *in situ* hybridisation

Fluorescent *in situ* hybridizations using digoxigenin-labelled *c-fos* were performed on dissected juvenile zebrafish with few modification to the original method (Brend & Holley, 2009). After overnight fixation in 4% PFA, protein K treatment (2mg/ml 20 minutes of incubation), inactivation of endogenous peroxidase with H2O2 (22%v/v for 30 min at room temperature), additional fixation (30 minutes at room temperature) and 3h of incubation with the hybridisation buffer, fish were incubated with the *c-fos* probe (courtesy from Ricardo Neto Silva (Forward CCGATACACTGCAAGCTGAA and Reverse ATTGCAGGGCTATGGAAGTG), or with DAT probe (Filippi et al., 2010), or Slc6a4b (Norton, Folchert, & Bally-Cuif, 2008). C*-fos, DAT* and *Slc6a4b* probes were detected with anti-Digoxigenin-POD, Fab fragments (Roche, 1:3000) and TSA Plus Cyanine 3 System (Perkin Elmer, 1:50). Nuclear staining was obtained using DAPI (Sigma-Aldrich, 1: 500). Fish were than mounted for imaging in low melting point agarose (2.5% Agarose, 0.8% glycerol, PBS-Tween) and imaged.

### Imaging and registration

A custom built two-photon microscope (INSS) was used for image acquisition of whole-brain *in situ*s. Both DAPI and Cy3 Images were collected with a 10x objective (Olympus, W Plan-Apochromat 10x/0.5 M27 75mm) using a “Chameleon” titanium–sapphire laser tuned to 1030 nm (Coherent Inc., Santa Clara, CA, U.S.A.) and controlled using custom written software in LabView.. Registration of *in-situ* images was performed using ANTs version 2.1.0 (Advanced Normalisation Tools, https://www.ncbi.nlm.nih.gov/pubmed/17659998) running on the UCL Legion compute cluster. Images were down-sampled to 512*512 and parameters were slightly modified from Marquart et al., 2017 fixed registration:

*antsRegistration -d 3 --float 1 -o [Registered_Image_, Registered_Image _warped*.*nii*.*gz] --interpolation WelchWindowedSinc --use-histogram-matching 0 -r [reference_Image, Registered_Image,1] -t rigid[0*.*1] - m MI[reference_Image, Registered_Image _0*.*nii,1,32,Regular,0*.*25] -c [1000×500×250×100,1e-8,10] -- shrink-factors 12×8×4×2 -s 4×3×2×1 -t Affine[0*.*1] -m MI[reference_Image, Registered_Image,1,32,Regular,0*.*25] -c [1000×500×250×100,1e-8,10] --shrink-factors 12×8×4×2 -s 4×3×2×1 -t SyN[0*.*1,6,0] -m CC[reference_Image, Registered_Image _0*.*nii,1,2] -c [1000×500×500×250×100,1e-7,10] --shrink-factors 12×8×4×2×1 -s 4×3×2×1×0*

*antsApplyTransforms -d 3 -v 0 --float -n WelchWindowedSinc -i Registered_Image _1*.*nii -r reference_Image -o Registered_Image _warped_red*.*nii*.*gz -t Registered_Image _1Warp*.*nii*.*gz -t Registered_Image _0GenericAffine*.*mat*

The registered image stacks were then normalised to adjust for intensity variations between imaging sessions. caused by a variety of sources (staining efficiency, laser power fluctuations, light detector sensitivity, etc.). Normalization was accomplished by computing an intensity histogram for each fish brain’s volume (with 10000 discrete intensity bins spanning the range -4000.0 to 70000.0) for all 512*512*273 voxels. The minimum value bin (with at least 100 voxels) was used as the bias offset, and subtracted from all voxel values. The mode value, minus the bias, provided a robust estimate of the background/baseline fluorescence and was thus used to normalize voxel values for the entire volume. Therefore, after normalization, intensity value of 1 reflected the background level, 2 reports fluorescence level that is twice the background, and so on. Histogram normalization was performed for each individual fish brain volume prior to any region or voxel-based analysis.

**Figure 2B and 3A** Reconstruction of cross section images were obtained by using the Fiji “Volume viewer” plugin. Schematics of cross- and horizontal-section were obtained by using the “Neuroanatomy of the zebrafish brain”

**Figure 2D and 3B** Percentages of *c-fos* activation were calculated for each of the 6 different areas highlighted in Figure 2B and 3A, using custom written Python functions, in the following way. A 3D mask for each area was generated by using the “Segmentation Editor” plugin Fiji (https://imagej.net/Segmentation_Editor). *C-fos* percentage values for each condition (C (+S), C (-S), Fi (-S), Pi (-S)) were obtained by subtracting and then dividing each *c-fos* average value of the mask by the basal *c-fos* average value calculated in control fish No Social Cue.

### Statistics

Statistical analysis was performed using Python scipy stats libraries. Since VPI, percent time moving, and *c-fos* activity distributions were generally not normally distributed, we used the non-parametric Mann-Whitney U-test of independent samples for hypothesis testing throughout the manuscript.

### Drug treatment

Juvenile fish were treated with 30µM or 50µM Buspirone (Buspirone HCl, Sigma) for 15 minutes prior the experiment. After washing, fish were run through the behavioural assay. Each fish was used only once.

## Data availability

All the images, video, protocols, analysis scripts, and data that support the findings of this study are available either from this website (http://www.dreo-sci.com/resources/) or from the corresponding author upon request.

## Acknowledgements

The authors acknowledge Steve Wilson for providing lab resources. Ricardo Neto Silva for the *c-fos* probe. Jason Rihel for some of the reagents. Wolfgang Driver for the dopaminergic probes and William Norton for the serotoninergic probes. UCL fish facility team for fish care/husbandry. This work was supported by the Wellcome Trust Grant Ref. 202465/Z/16/Z.

## Author contributions

H.T. designed the experiments, performed the experiments, analyzed the data, helped to write the manuscript. M.V-P. performed the serotonin and dopaminergic experiments. T.R. helped with the analysis and writing the paper. A.R.K. designed the software for the image analysis after registration and helped to write the manuscript. E.D. conceived the project, designed the experiments, performed some of the experiments and data analysis, and helped write the manuscript.

## References

Barba-Escobedo, P. A., & Gould, G. G. (2012). Visual social preferences of lone zebrafish in a novel environment: Strain and anxiolytic effects. Genes, Brain and Behavior, 11(3), 366–373. https://doi.org/10.1111/j.1601-183X.2012.00770.x

Bencan, Z., Sledge, D., & Levin, E. D. (2009). Buspirone, chlordiazepoxide and diazepam effects in a zebrafish model of anxiety. Pharmacology Biochemistry and Behavior, 94(1), 75–80. https://doi.org/10.1016/j.pbb.2009.07.009

Brend, T., & Holley, S. A. (2009). Zebrafish whole mount high-resolution double fluorescent in situ hybridization. J Vis Exp, (25). https://doi.org/10.3791/1229

Cacioppo, J T, Cacioppo, S., Capitanio, J. P., & Cole, S. W. (2015). The Neuroendocrinology of Social Isolation. Annual Review of Psychology, Vol 66, 66, 733–767. https://doi.org/10.1146/annurev-psych-010814-015240

Cacioppo, John T., Norris, C. J., Decety, J., Monteleone, G., & Nusbaum, H. (2009). In the eye of the beholder: Individual differences in perceived social isolation predict regional brain activation to social stimuli. Journal of Cognitive Neuroscience, 21(1), 83–92. https://doi.org/10.1162/jocn.2009.21007

Dreosti, E., Lopes, G., Kampff, A. R., & Wilson, S. W. (2015). Development of social behavior in young zebrafish. Frontiers in Neural Circuits, 9. https://doi.org/ARTN3910.3389/fncir.2015.00039

Egan, R. J., Bergner, C. L., Hart, P. C., Cachat, J. M., Canavello, P. R., Elegante, M. F., … Kalueff, A. V. (2009). Understanding behavioral and physiological phenotypes of stress and anxiety in zebrafish. Behavioural Brain Research, 205(1), 38–44. https://doi.org/10.1016/j.bbr.2009.06.022

Engeszer, R. E., Ryan, M. J., & Parichy, D. M. (2004). Learned social preference in zebrafish. Current Biology, 14(10), 881–884. https://doi.org/DOI10.1016/j.cub.2004.04.042

File, S. E., & Seth, P. (2003, February 28). A review of 25 years of the social interaction test. European Journal of Pharmacology, Vol. 463, pp. 35–53. https://doi.org/10.1016/S0014-2999(03)01273-1

Filippi, A., Mahler, J., Schweitzer, J., & Driever, W. (2010). Expression of the Paralogous Tyrosine Hydroxylase Encoding Genes th1 and th2 Reveals the Full Complement of Dopaminergic and Noradrenergic Neurons in Zebrafish Larval and Juvenile Brain. Journal of Comparative Neurology, 518(4), 423–438. https://doi.org/10.1002/cne.22213

Giacomini, A. C. V. V, de Abreu, M. S., Koakoski, G., Idalencio, R., Kalichak, F., Oliveira, T. A., … Barcellos, L. J. G. (2015). My stress, our stress: Blunted cortisol response to stress in isolated housed zebrafish. Physiology & Behavior, 139, 182–187. https://doi.org/10.1016/j.physbeh.2014.11.035

Gould, G. G., Hensler, J. G., Burke, T. F., Benno, R. H., Onaivi, E. S., & Daws, L. C. (2011). Density and function of central serotonin (5-HT) transporters, 5-HT 1A and 5-HT2A receptors, and effects of their targeting on BTBR T+tf/J mouse social behavior. Journal of Neurochemistry, 116(2), 291–303. https://doi.org/10.1111/j.1471-4159.2010.07104.x

Herrera, D. G., & Robertson, H. A. (1996). Activation of c-fos in the brain. Prog Neurobiol, 50(2–3), 83–107. https://doi.org/10.1016/s0301-0082(96)00021-4

Huang, C. C., Lu, R. B., Yen, C. H., Yeh, Y. W., Chou, H. W., Kuo, S. C., … Huang, S. Y. (2015). Dopamine transporter gene may be associated with bipolar disorder and its personality traits. European Archives of Psychiatry and Clinical Neuroscience, 265(4), 281–290. https://doi.org/10.1007/s00406-014-0570-0

Lalonde, R., & Strazielle, C. (2009). Relations between open-field, elevated plus-maze, and emergence tests in C57BL/6J and BALB/c mice injected with GABA- and 5HT-anxiolytic agents. Fundamental & Clinical Pharmacology, 24(3), 365–376. https://doi.org/10.1111/j.1472-8206.2009.00772.x

Lau, B. Y. B., Mathur, P., Gould, G. G., & Guo, S. (2011). Identification of a brain center whose activity discriminates a choice behavior in zebrafish. Proceedings of the National Academy of Sciences of the United States of America, 108(6), 2581–2586. https://doi.org/10.1073/pnas.1018275108

Lillesaar, C. (2011). The serotonergic system in fish. J Chem Neuroanat, 41(4), 294–308. https://doi.org/10.1016/j.jchemneu.2011.05.009

Marquart, G. D., Tabor, K. M., Horstick, E. J., Brown, M., Geoca, A. K., Polys, N. F., … Burgess, H. A. (2017). High-precision registration between zebrafish brain atlases using symmetric diffeomorphic normalization. Gigascience, 6(8). https://doi.org/10.1093/gigascience/gix056

McDowell, A. L., Dixon, L. J., Houchins, J. D., & Bilotta, J. (2004). Visual processing of the zebrafish optic tectum before and after optic nerve damage. Visual Neuroscience, 21(2), 97–106. https://doi.org/10.1017/S0952523804043019

Norton, W. H. J., Folchert, A., & Bally-Cuif, L. (2008). Comparative analysis of serotonin receptor (HTR1A/HTR1B families) and transporter (slc6a4a/b) gene expression in the zebrafish brain. Journal of Comparative Neurology. https://doi.org/10.1002/cne.21831

O’Connell, L. A., & Hofmann, H. A. (2011). The vertebrate mesolimbic reward system and social behavior network: a comparative synthesis. J Comp Neurol, 519(18), 3599–3639. https://doi.org/10.1002/cne.22735

Patel, S., & Hillard, C. J. (2006). Pharmacological evaluation of cannabinoid receptor ligands in a mouse model of anxiety: Further evidence for an anxiolytic role for endogenous cannabinoid signaling. Journal of Pharmacology and Experimental Therapeutics, 318(1), 304–311. https://doi.org/10.1124/jpet.106.101287

Rogers-Carter, M. M., Varela, J. A., Gribbons, K. B., Pierce, A. F., McGoey, M. T., Ritchey, M., & Christianson, J. P. (2018). Insular cortex mediates approach and avoidance responses to social affective stimuli. Nat Neurosci, 21(3), 404–414. https://doi.org/10.1038/s41593-018-0071-y

Schneier, F. R., Saoud, J. B., Campeas, R., Fallon, B. A., Hollander, E., Coplan, J., & Liebowitz, M. R. (1993). Buspirone in social phobia. Journal of Clinical Psychopharmacology, 13(4), 251–256. https://doi.org/10.1097/00004714-199308000-00004

Shams, S, Amlani, S., Buske, C., Chatterjee, D., & Gerlai, R. (2018). Developmental social isolation affects adult behavior, social interaction, and dopamine metabolite levels in zebrafish. Dev Psychobiol, 60(1), 43–56. https://doi.org/10.1002/dev.21581

Shams, S, Seguin, D., Facciol, A., Chatterjee, D., & Gerlai, R. (2017). Effect of social isolation on anxiety-related behaviors, cortisol, and monoamines in adult zebrafish. Behav Neurosci, 131(6), 492–504. https://doi.org/10.1037/bne0000220

Shams, Soaleha, Amlani, S., Buske, C., Chatterjee, D., & Gerlai, R. (2018). Developmental social isolation affects adult behavior, social interaction, and dopamine metabolite levels in zebrafish. Developmental Psychobiology, 60(1), 43–56. https://doi.org/10.1002/dev.21581

Sloan Wilson, D., Clark, A. B., Coleman, K., & Dearstyne, T. (1994). Shyness and boldness in humans and other animals. Trends Ecol Evol, 9(11), 442–446. https://doi.org/10.1016/0169-5347(94)90134-1

van Vliet, I. M., den Boer, J. A., Westenberg, H. G., & Pian, K. L. (1997). Clinical effects of buspirone in social phobia: a double-blind placebo-controlled study. The Journal of Clinical Psychiatry, 58(4), 164–168. https://doi.org/10.4088/jcp.v58n0405

Winslow, J. T. (2003). Mouse social recognition and preference. Curr Protoc Neurosci, Chapter 8, Unit 8 16. https://doi.org/10.1002/0471142301.ns0816s22

Zellner, D., Padnos, B., Hunter, D. L., MacPhail, R. C., & Padilla, S. (2011). Rearing conditions differentially affect the locomotor behavior of larval zebrafish, but not their response to valproate-induced developmental neurotoxicity. Neurotoxicol Teratol, 33(6), 674–679. https://doi.org/10.1016/j.ntt.2011.06.007

Ziv, L., Muto, A., Schoonheim, P. J., Meijsing, S. H., Strasser, D., Ingraham, H. A., … Baier, H. (2013). An affective disorder in zebrafish with mutation of the glucocorticoid receptor. Molecular Psychiatry, 18(6), 681–691. https://doi.org/10.1038/mp.2012.64

