## Supplementary Figure 1 for "The lonely fish is not a loner fish: whole-brain mapping reveals abnormal activity in socially isolated zebrafish"

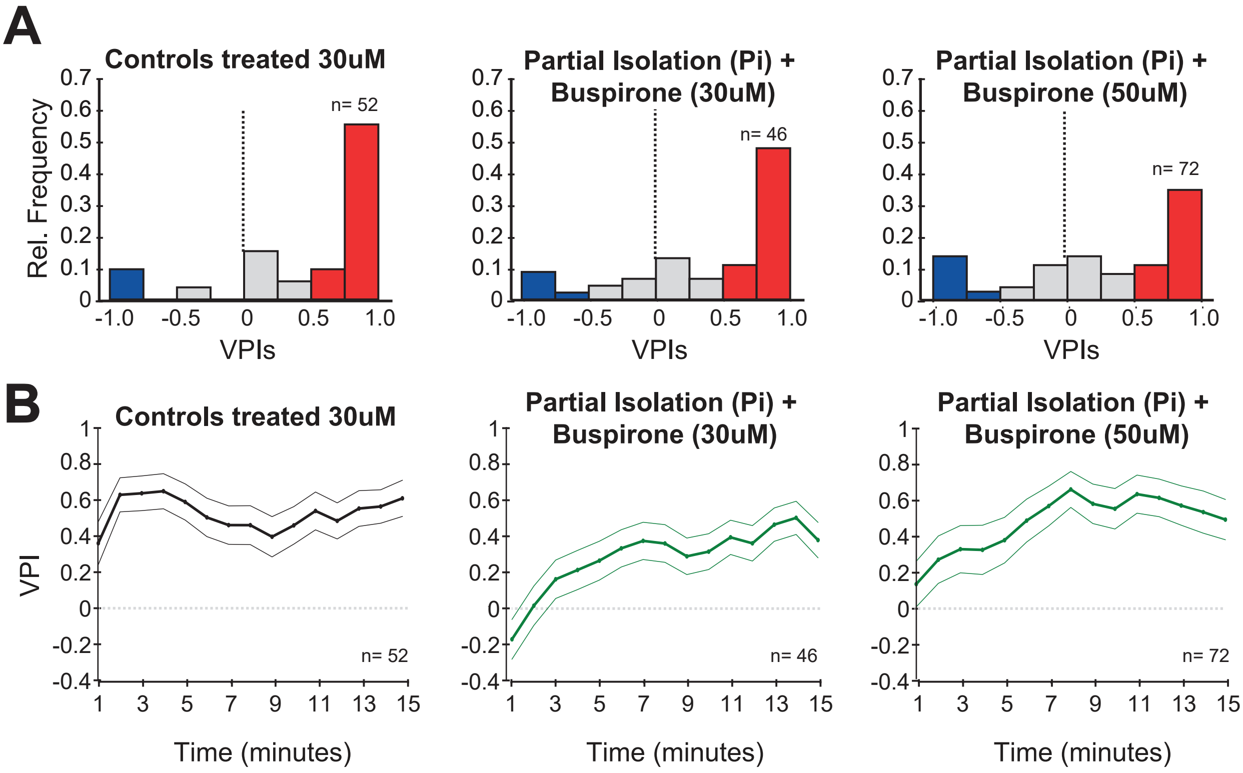


**Supplementary Figure 1. Buspirone rescues social preference in isolated fish.**

**A.** Histograms of VPIs during the social cue period in control (C) fish treated with 30 μM of Buspirone, in Partially isolated (Pi) fish treated with 30 μM and 50 μM of Buspirone. or visual clarity, the bars are colored as in Figure 1. **B.** VPI values calculated in one minute time bins for controls treated with 30μM of Buspirone, Pi fish treated with 30μM, and Pi fish treated with 50μM of Buspirone.
