## Supplementary Movie Legend for "The lonely fish is not a loner fish: whole-brain mapping reveals abnormal activity in socially isolated zebrafish"

### Supplementary -Text for Avi File
#### Movie S1.

**Example of a control and a fully isolated +S fish video during social cue presentation.** Two minutes of behaviour is shown in 20 seconds (6x playback acceleration). The control fish shows a strong social preference for the social cue and has a stereotypical social phenotype (left). The test fish spends most of its time watching the social cue with a 45-degree angle and synchronizing its bout motion with the other two conspecifics. The fully isolated fish spends long periods of time as well on the side of the conspecifics. Its behaviour, however, its characterized by long pauses while watching the conspecifics (right).
